# Signaling via GABA_B_ receptors regulates early development and neurogenesis in the basal metazoan *Nematostella vectensis*

**DOI:** 10.1101/621060

**Authors:** Shani Levy, Vera Brekhman, Anna Bakhman, Arnau Sebé-Pedrós, Mickey Kosloff, Tamar Lotan

**Author notes:** Corresponding authors: TL -; MK.

## Abstract

The metabotropic gamma-amino-butyric acid B receptor (GABA_B_R) is a G protein–coupled receptor that mediates neuronal inhibition by the neurotransmitter GABA. Here, we identified putative GABA_B_ receptors and signaling modulators in the basal sea anemone *Nematostella vectensis*. Activation of GABA_B_R signaling reversibly arrests planula-to-polyp transformation during early development and affects the neurogenic program. We identified four *Nematostella* GABA_B_R homologs that have the conserved 3D extracellular domains and residues needed for binding of GABA and the GABA_B_R agonist baclofen. Transcriptomic analysis, combined with spatial analysis of baclofen-treated planulae, revealed that baclofen down-regulated pro-neural factors such as *NvSoxB(2)*, *NvNeuroD1* and *NvElav1*. Baclofen also inhibited neuron development and extended neurites, resulting in an under-developed and less organized nervous system. Our results shed light on cnidarian development and suggest an evolutionarily conserved function for GABA_B_R in regulation of neurogenesis, highlighting *Nematostella* as a new model system to study GABA_B_R signaling.

## Introduction

γ-aminobutyric acid (GABA) is the major inhibitory neurotransmitter in both vertebrates and invertebrates, playing a key role in modulating neuronal activity. GABA activity is mediated by two distinct receptors – the ionotropic GABA_A_ and the metabotropic GABA_B_ receptors. The GABA_A_ receptor is a chloride ion channel that mediates fast synaptic inhibition. The GABA_B_ receptor (GABA_B_R) is a G-protein-coupled receptor (GPCR) that produces prolonged and slow synaptic inhibition through second messengers and modulation of calcium (Ca^2+^) and potassium (K^+^) channels. These receptors function as obligatory heterodimers composed of GABA_B1_ and GABA_B2_ subunits, with both subunits required for their function (1–3). GABA binds to a large Venus flytrap (VFT) structural module in the extracellular portion of the GABA_B1_ monomer (4, 5), whereas GABA_B2_ does not bind GABA, but activates the coupled G protein and enhances ligand affinity via interactions with the GABA_B1_ monomer (6–10). The VFT module is not unique to GABA_B1_R and can be found in diverse proteins, ranging from other GPCRs and ion channels in eukaryotes and periplasmic binding proteins in prokaryotes (11). During embryonic development, GABA_B_R plays important roles in mammalian neuronal proliferation, migration and network formation (12–15). In adults, in addition to a general role in inhibiting neurotransmission, GABA_B_R signaling is also an important inhibitor of neuronal differentiation, controlling stem and progenitor cell proliferation (16, 17).

Although GABA_B_ receptors have been extensively studied in mammalian model organisms and in *Drosophila* and *C. elegans* (18–21), little is known on their function in non-bilaterian animals. Cnidaria are the phylogenetic sister group to Bilateria and one of the earliest-branching metazoan taxa that possess a nervous system (22). Other early metazoans, such as Placozoa and Porifera, possess genes related to sensory transmission, yet lack neurons and synapses, while Ctenophora harbor a nervous system of unclear homology to cnidarian and bilaterian systems (23–25). Moreover, this lineage is missing most of the Bilateria classical neurotransmitters, including GABA (26). By contrast, cnidarians possess a diffuse nervous system that contains three primary types of neuronal cells: sensory neurons, ganglion neurons, and cnidocytes, the stinging cells that characterize the phylum (27, 28). Their phylogenetic placement and simple nerve-net structure make them an attractive model for exploring basic neurogenic processes.

Among the cnidarians, the sea anemone *Nematostella vectensis* has become an important model system, with a published genome and available genetic tools (29, 30). During its simple life cycle, an embryo develops into a larval stage of a swimming planula, which metamorphoses into a primary polyp that eventually gives rise to the mature polyp (31). Despite its relatively simple bodyplan, analysis of the *Nematostella* genome has revealed an unexpected complexity and extensive conservation of vertebrate genomic content and organization (30). This similarity is also reflected in a conserved neuronal gene repertoire, which includes orthologous genes associated with cholinergic, glutamatergic, and aminergic neurotransmission (27, 32-34). Genes putatively related to GABA signaling were identified in *Nematostella* using large-scale phylogenetic analysis (32). GABA itself was shown to accumulate early in *Nematostella* planulae in the aboral pole and in ectodermal neurons (34). However, the role of GABA during development or in neurogenesis is unknown and the specific receptors that mediate GABA signaling have not been characterized.

In this study, we demonstrated that *Nematostell*a has four putative GABA_B1_R genes. The extracellular domains of these genes are predicted to have a homologous 3D structure to mammalian GABA_B1_R and contain conserved residues that can mediate agonist binding. Using baclofen, a GABA_B_R-specific agonist, we elucidated the role of GABA-mediated signaling during *Nematostell*a early development. Our findings showed that baclofen arrested planula-to-polyp transformation and inhibited neurogenesis; yet, the effect was reversible. These results suggest a conserved role for GABA signaling in controlling neuronal network formation in the basal sea anemone and highlights future uses for *Nematostell*a as a model organism to study GABA-mediated signaling.

## Results

### GABA_B_ receptors play a role in planula-to-polyp transformation

The morphogenetic transition of *Nematostella* planulae into primary polyps occurs 7 to 12 days post-fertilization (dpf). Since GABA was found to accumulate in early *Nematostella* planulae (34), we tested the effect of GABA on planula-to-polyp transformation. We found that addition of GABA (10^−3^ M) at 3 dpf, at the early planula stage, prevented metamorphosis of more than 85% of the planulae, while about 80% of control planulae metamorphosed by 8 dpf (Fig. 1a). A lower GABA concentration (10^−4^ M) reduced the rate of metamorphosis, but did not inhibit the process, probably due to natural GABA degradation (Fig. 1a). Arrested planulae continued to develop to the late planula stage, although their development was much slower than untreated control planulae (Fig. 1b). To identify which GABA receptor family is involved in the inhibition of transformation, we tested the effect of GABA_A_R and GABA_B_R agonists. We found that the GABA_A_R agonist muscimol (10^−4^ M) demonstrated toxic and lethal effects, inhibiting further development of early planulae (data not shown). By contrast, the GABA_B1_R agonist baclofen (10^−4^ M) had a similar physiological effect as GABA, but inhibited metamorphosis to a larger degree (Fig. 1a). Additionally, CGP-7930, a GABA_B2_R positive allosteric modulator (35), prevented metamorphosis in more than 60% of the planulae and the affected planulae had a similar morphology as GABA- or baclofen-treated planulae (Fig. 1b). Baclofen-treated planulae did not transform to primary polyps, but their body structure, marked by F-actin, displayed a similar pattern to control untreated planulae (Supplementary Fig. 1). However, in control oral poles, tentacles emerged and elongated around the oral opening, whereas in GABA/baclofen/CGP-7930-treated planulae tentacle growth was arrested (Fig. 1b).

**Figure 1:**
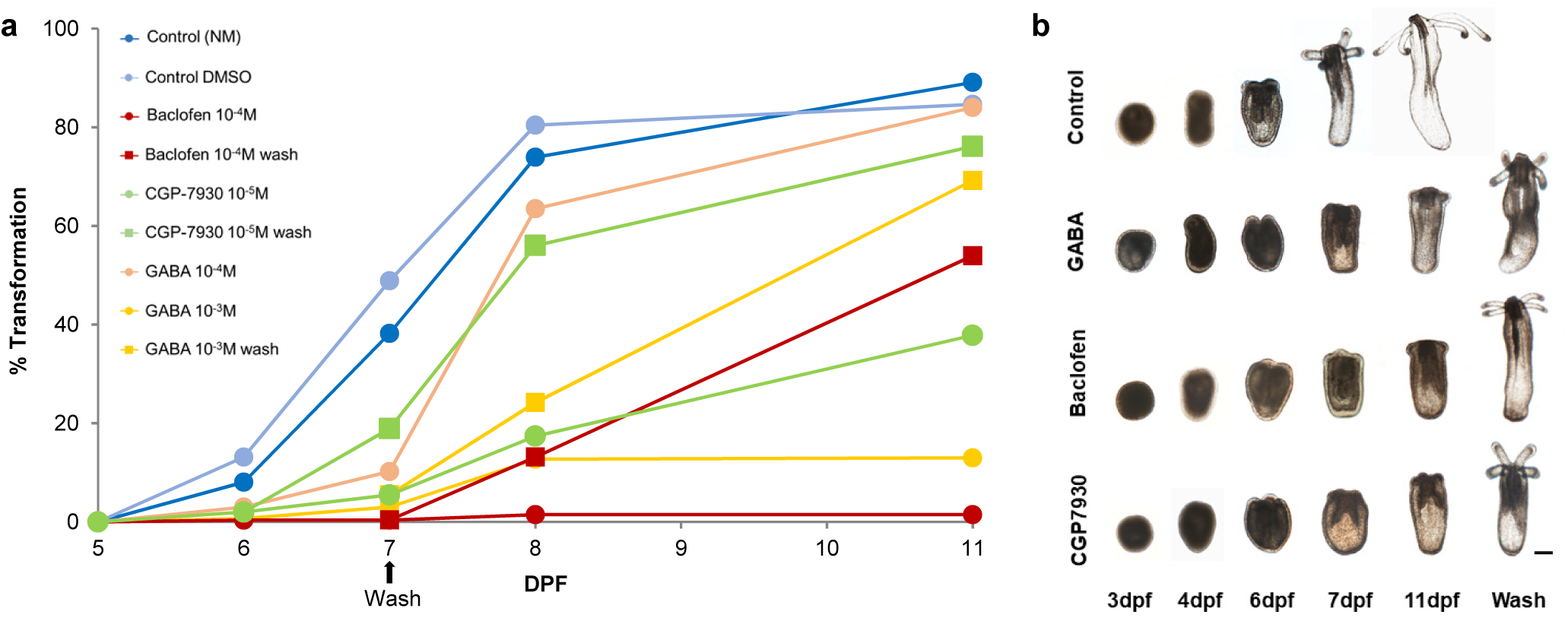
GABA and specific GABA_B_R agonist and modulator cause reversible inhibition of *Nematostella* metamorphosis. **a,** Percentage of planulae that transformed to polyps as a function of time (dpf) in controls (NM, DMSO) and in planulae treated with GABA (10^−3^M, 10^−4^M), the GABA_B1_R agonist baclofen (10^−4^M), and the positive GABA_B2_R allosteric modulator CGP-7930 (10^−4^M). Controls and agonists (circles) were added at 3 dpf. At 7 dpf, agonists were removed in half of the samples (rectangles). **b,** Planula development and metamorphosis under control conditions and upon GABA, baclofen or CGP-7930 treatment. Note that treated planulae were smaller and failed to elongate or metamorphose. Four days after removal of compounds, planulae recovered and transformed into primary polyps (wash, 11 dpf). Scale bar, 100 µm.

Because oralization and tentacle induction were previously shown to be controlled by the Wnt pathway (36, 37), we tested the expression of specific Wnt ligands that were shown to characterize the oral pole of *Nematostella* planulae (38). We found that the expression levels and oral localization of *NvWntA*, *NvWnt1*, *NvWnt3*, *NvWnt5* and *NvWnt11* did not change following baclofen treatment (Supplementary Fig. 2) suggesting that the inhibitory effect of GABA_B_R signalling is not mediated through changes in the expression of Wnt ligands.

In addition, planulae treated with baclofen exhibited similar apical tufts as control planulae (Supplementary Fig. 1), an organ that was suggested to play a role during *Nematostella* metamorphosis (39, 40). Yet, in control planulae the apical tuft was lost during metamorphosis, whereas in baclofen-treated planulae it remained. The effects of GABA, baclofen and CGP-7930 were reversible, as removal of these compounds on 7 dpf restored metamorphosis and enabled the planulae to develop into primary polyps (Fig. 1a). However, the addition of GABA_B_R antagonists such as saclofen, phaclofen and CGP-54626 had no detectable effect on planulae development or metamorphosis rates (data not shown). These results suggest that a unique GABA_B_R homolog, which is resistant to known antagonists, mediates the effect of GABA on planula-to-polyp transformation.

### Identification and characterization of *Nematostella* GABA_B_ receptors

To identify *Nematostella* GABA_B_ receptors, we searched all *Nematostella* proteins in the NCBI RefSeq database using the sequences of human GABA_B1_ and GABA_B2_ receptors as queries – both with the full length proteins and with the N-terminal extracellular domain and the transmembrane regions separately. Putative *Nematostella* GABA_B_R homologs were used as queries against the Uniprot database to confirm homology; indeed, the top hits for all identified genes were GABA_B_R genes. This search identified eight candidates GABA_B_R homologs in *Nematostella*. Seven of these genes appeared in a large genomic survey of chemical transmission-related genes (32): four genes (v1g244104, v1g239821, v1g206093, v1g243252) had full sequences, but three other homologs had only partial sequences (see Methods for details). We, therefore, fully sequenced the four partial or new genes, which were assigned the following NCBI identifiers: MH194577-79 and MH355581.

The GABA_B_R homologs we identified contained the conserved domains expected of a functional GABA_B_ receptor (Fig. 2). These included a signal peptide, followed by a large extracellular N-terminal domain that contains the structural VFT module that binds GABA, as well as seven trans-membrane (TM) domains that are characteristic of G protein-coupled receptors. Predicted coiled-coil motifs, which are found in the human GABA_B_R where they mediate hetero-dimerization and receptor trafficking (41, 42), were predicted in only three *Nematostella* homologs. In general, the intracellular C-terminus portion of the *Nematostella* homologs had low similarity to the corresponding human sequences.

**Figure 2:**
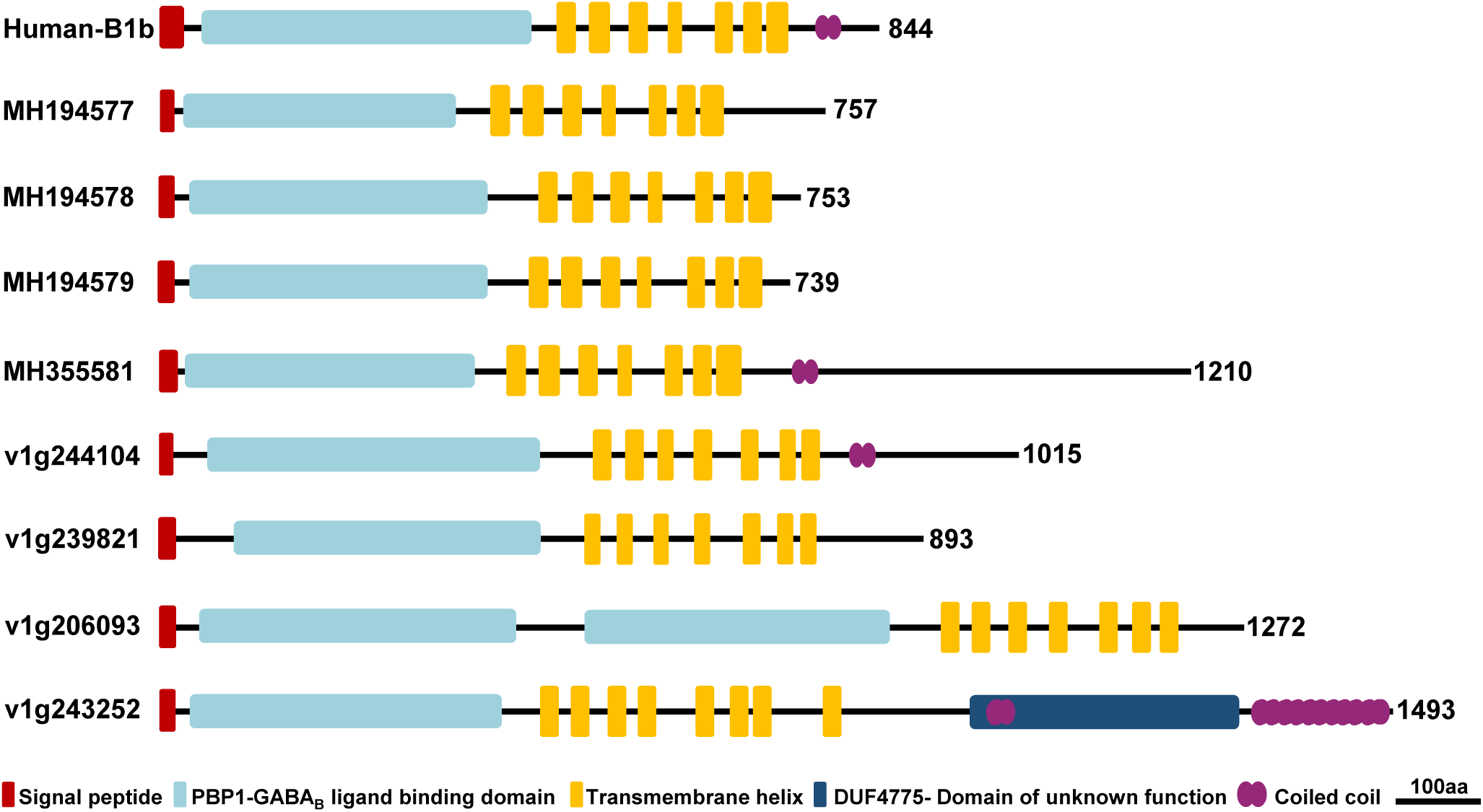
Schematic representation of predicted domains in putative *Nematostella* GABA_B_R homologs in comparison to human GABA_B1_R. The eight *Nematostella* proteins contain a conserved signal peptide, an extracellular “Periplasmic Binding Protein type1 (PBP1) GABA_B_ ligand binding domain” (the structural VFT module that in mammals forms binds GABA), predicted helical transmembrane domains, and coiled-coil domains. The domains were predicted as detailed in the Materials and Methods.

Six of these homologs had the same global domain arrangement as human GABA_B_R. One *Nematostella* homolog (v1g206093) contained two extracellular domains, each corresponding to a separate predicted VFT module. These two domains are 26% identical in sequence, suggesting that they have dissimilar function (Supplementary Fig. 3). Another homolog, v1g243252, had eight predicted TM helices (predicted by three different prediction servers, see Materials and Methods) and a ∼300 residue domain of unknown function (DUF4475) located after these TM domains.

To test if the identified GABA_B_R homologs exhibit neuronal-related expression, we searched the recently published *Nematostella* cell type expression atlas (43), quantifying the expression of the eight GABA_B_R homologs (Supplementary Fig. 4). We detected all eight homologs in this single cell analysis, although MH194578 and MH194579 were expressed at extremely low levels and did not show any cell type specificity, while MH355581 was expressed at low levels and only in a specific group of cells of the gastrodermis. In contrast, four of the GABA_B_R homologs (MH194577, v1g244104, v1g239821, and v1g206093) were expressed in neuronal cell clusters, including in larval neurons. On the other hand, v1g243252 was identified only in adult digestive filaments – this non-neural expression, together with its divergent protein architecture, suggested it is not a GABA_B_R homolog that can mediate the phenotypes observed above.

Unlike many GPCRs, the GABA binding site is located exclusively in the extracellular VFT domain of the GABA_B1_R. However, the VFT module can be found in many proteins that do not bind GABA (11). Therefore, to assess the ability of these candidate homologs to act as functional GABA-binding receptors, we compared their extracellular domains to their bilaterian homologs. Using multiple sequence alignments and the 3D structures of human GABA_B1_R, we analyzed the conservation of residues shown previously as critical for ligand binding. This comparison revealed that only four sequences had both the homologous scaffold structure and the conserved residues needed for agonist binding, while the other four sequences were either missing essential structural and binding residues or had an insertion that was predicted to interfere with agonist binding (Fig. 3 and Supplementary Fig. 3).

**Figure 3:**
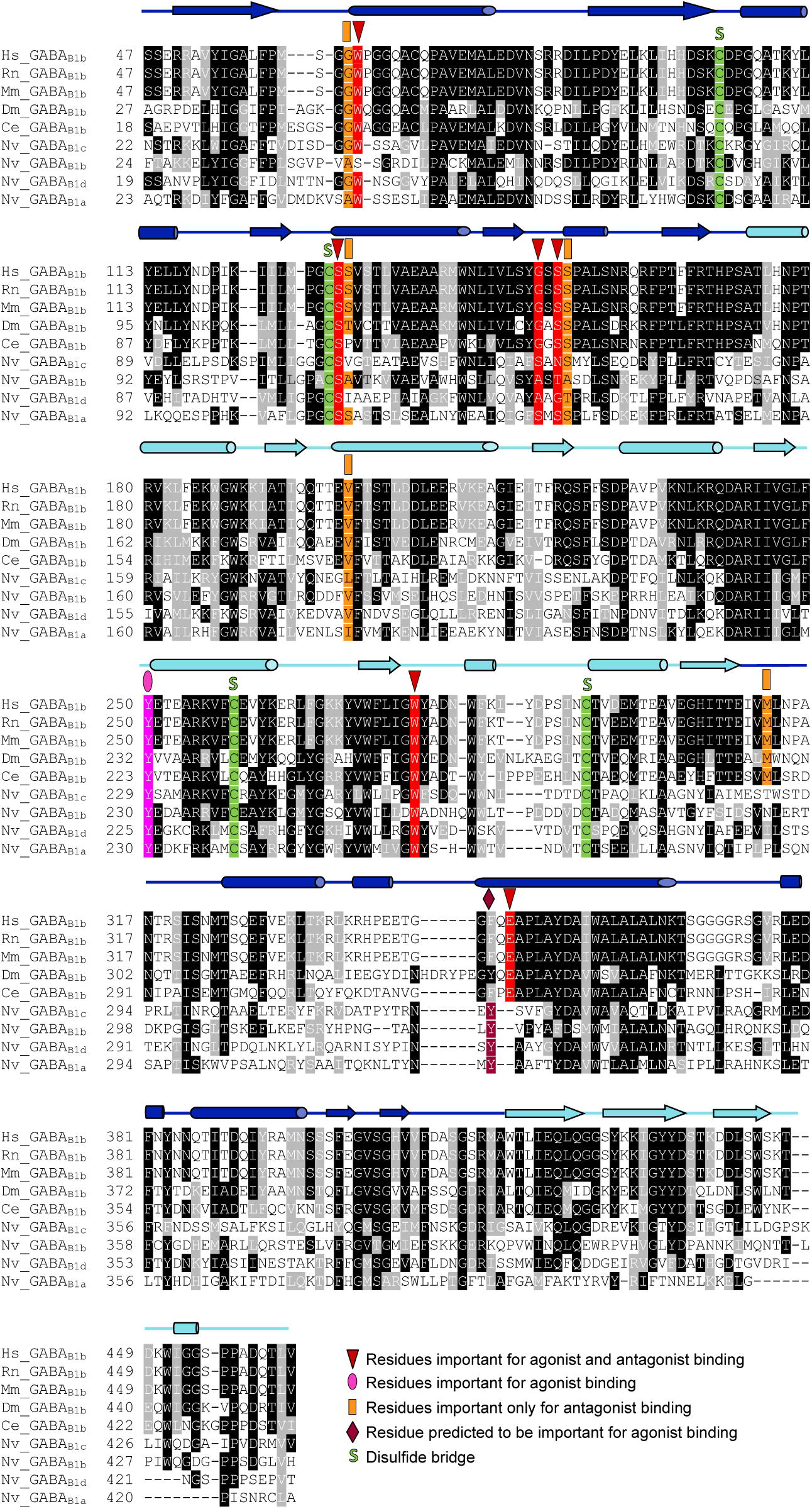
Sequence comparison of the extracellular regions in bilaterian and putative *Nematostella* GABA_B1_R homologs. Identical residues are marked in black and conservatively substituted residues are marked in gray. Residues involved in both agonist and antagonist binding are marked in red, residues involved in agonist binding only are marked in magenta, residues predicted to be involved in agonist binding are marked in bordeaux and residues involved in antagonist binding only are marked in orange. Cysteines showed to form disulfide bridges (S) that stabilize the VFT domain and are critical for GABA binding are marked in green. Secondary structure elements of human GABA_B1_R are displayed above the alignment: α-helices are shown as cylinders and β-strands as arrows, colored purple and cyan in the LB1 and LB2 sub-domains of the extracellular VFT domain, respectively. Bilaterian sequences accession numbers: human (Hs), NP_068703; rat (Rn), AAD19657; mouse (Mn), AAH56990; *Drosophila* (DM), AAF53431; *C. elegans* (Ce), ACE63490 and *Nematostella* (Nv) sequences: Nv_GABA_B1a_, MH194577; Nv_GABA_B1b_, MH194578; Nv_GABA_B1c_, MH194579; Nv_GABA_B1d_, MH355581.

Initially, we marked which conserved residues determine a 3D scaffold for ligand binding in vertebrate GABA_B1_R. The crystal structure of the VFT extracellular domain of human GABA_B1_R consists of two lobe-shaped domains (LB1 and LB2) that close upon ligand binding. This structure is stabilized by two conserved disulfide bridges, Cys103-Cys129 and Cys259-Cys293 (Fig. 3), which were shown to be essential for a functional GABA-binding structure in the GABA_B1_R (5). Five *Nematostella* proteins (MH194577-79, MH355581 and v1g244104) contain four cysteines that correspond to all of these disulfide bridge-forming residues, whereas the other three homologs (v1g239821, v1g206093, and v1g243252) were missing cysteines that correspond to Cys103 and Cys129 – these three proteins are therefore predicted not to bind GABA (Fig. 3 and Supplementary Fig. 3).

In the human GABA_B1_R, Trp65, Ser130, Ser153, Tyr250 and Gly151 were identified previously as critical mediators of GABA binding (4, 5). Among the five *Nematostella* proteins, MH194577-79 and MH355581 contain identical residues or conserved substitutions in the corresponding positions, whereas v1g244104 had a large and positively-charged arginine in the position that corresponds to Ser153. Such a dramatic substitution will interfere with agonist binding via both steric and electrostatic interactions, suggesting that v1g244104 will not bind GABA or its analogs. In addition, the γ-amino group of agonists such as GABA or baclofen forms hydrogen bonds with Tyr250 in LB2 and His170 and Glu349 in LB1. All *Nematostella* GABA_B_R homologs contained a tyrosine residue in the corresponding position to Tyr250 and had a conserved tyrosine residue that the 3D modeling showed could substitute for Glu349 (Fig. 3). Since His170 was not conserved in any of the *Nematostella* GABA_B_ proteins, we suggest that His170 is dispensable for agonist binding. Taken together, these results predict that four *Nematostella* homologs, MH194577-79 and MH355581, contain functional GABA and baclofen binding sites and we therefore suggest that they are functional GABA_B1_R homologs.

We also analyzed the conservation of GABA_B1_R residues that were shown to mediate antagonist binding to human GABA_B1_R. Several human GABA_B1_R residues, such as Gly64, Ser131, Ser154, Val159, and Met 312, were suggested to mediate specific interactions with GABA_B1_R antagonists. While Gly64, Ser154 and Val159 were conserved in all *Nematostella* homologs, Ser131 was conserved only in MH194577 and was substituted by the similarly-sized alanine in MH355581. However, residues in *Nematostella* homologs that correspond to Met312 were not conserved, and the entire LB1 loop, in which Met 312 is located, was also not conserved (Fig. 3). Comparing the structure of the extracellular domain of human GABA_B1_R bound to the antagonist CGP54626 (4) with the predicted 3D structure of the corresponding domain in *Nematostella* showed that the first and the fifth loops in the predicted LB1 were longer by several amino acids than the corresponding loops in human GABA_B1_R (Fig. 4). These longer loops will most likely result in a substantially smaller binding pocket in the *Nematostella* homologs. Given that antagonists of GABA_B1_R are larger than the agonists of this receptor, we suggest that the smaller binding pocket of *Nematostella* homologs will prevent binding of these specific antagonists, explaining our experimental results.

**Figure 4:**
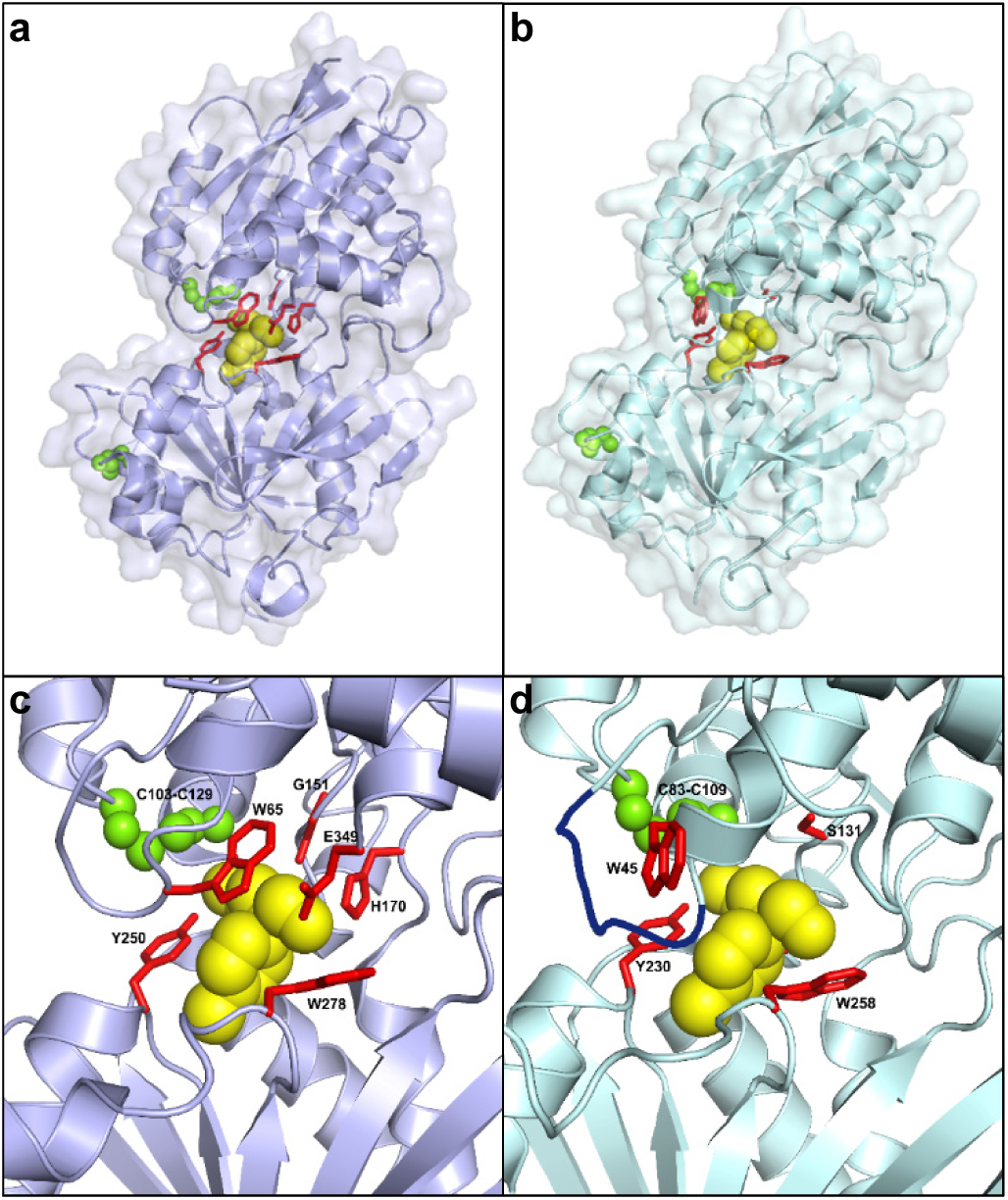
Three-dimensional visualization of the human and *Nematostella* GABA_B_R extracellular region bound to baclofen. **a**, The crystal structure of the extracellular VFT domain of the Human GABA_B1b_ receptor (PDB ID 4MS4). **b,** A model of the corresponding extracellular domain in *Nematostella* (GABA_B1a_ MH194577), built with Swiss-model using the human GABA_B1b_R structure (as in **a**) as a template. **c,** The GABA binding pocket in the human GABA_B1b_R. Residues that participate in agonist binding are shown as red sticks, baclofen is shown as yellow spheres, and disulfide bridges are shown as small green spheres. **d,** The predicted GABA binding pocket in *Nematostella* GABA_B1a_R. Conserved residues that correspond to GABA binding sites in the human GABA_B1b_R structure are shown as red sticks. Residues His170 and Glu349 in the human proteins do not have a corresponding residue in the *Nematostella* binding pocket. A longer loop in the *Nematostella* LB1 subdomain, which is predicted to prevent binding of human GABA_B1b_R antagonists, is colored dark blue.

### Transcriptome analysis

To identify genes and pathways involved in GABA_B_R signaling and in metamorphosis inhibition by the GABA agonist baclofen, we used a whole-transcriptome RNA-seq approach. Having observed no morphological difference in the inhibitory effects of baclofen at 3 dpf or 4 dpf, we analyzed samples from planulae at different time points from 4 dpf until 6 dpf, just before metamorphosis begins (Fig. 5a). Samples were analyzed at 2 h, 24 h and 48 h after baclofen addition. Because our metamorphosis experiments showed that the effect of baclofen on planulae was reversible (Fig. 1), we also sampled planulae 2 h and 24 h after baclofen removal. Control planulae were sampled at corresponding time points (Fig. 5a). On average, 15.77 million sequence reads were obtained for each sample. We continued with analysis of only differentially expressed transcripts (see Materials and Methods). Across all treatments, more than 10,000 transcripts displayed significant changes in gene expression (Supplementary Table 1). To examine the effect of the treatments over time, we explored the transcriptomic data using non-metric multidimensional scaling analysis (nMDS). This analysis showed clear differences between the transcriptomes of control and baclofen-treated planulae during the time of development (Fig. 5b). In addition, removal of baclofen dramatically affected the planula transcriptome within 2 h and, after additional 24 h, the transcriptome of washed planulae differed further from those of both baclofen-treated and control planulae (Fig. 5b). To demonstrate the effect of baclofen treatment together with changes in planulae developmental, we compared expression in different time points using an MA plot analysis (Fig. 5c,d). This revealed transcripts that changed significantly during development (at 24 h and 48 h) in both control and baclofen-treated planulae, as well as transcripts that significantly changed only with baclofen. Therefore, baclofen treatment did not arrest the planulae developmental plan, but had a specific affect that resulted in planulae-to-polyp transformation.

**Figure 5:**
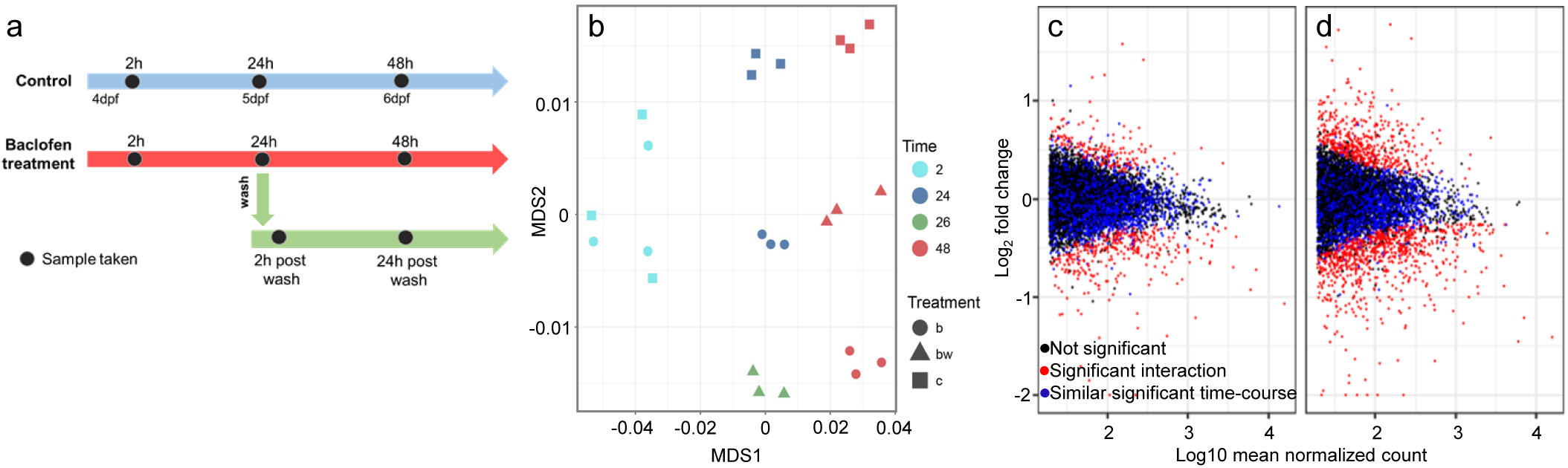
Transcriptomic analysis of control and baclofen-treated planulae. **a,** Schematic description of the experiment. Triplicate samples of control, baclofen-treated and baclofen-treated/washed planulae were taken at 4-6 dpf, 2 h, 24 h or 48 h post baclofen addition. 24 h after baclofen addition, half of the treated planulae were washed and samples were taken 2 h and 24 h post wash. **b,** nMDS analysis demonstrating the effect of baclofen (b) and baclofen wash (bw) on planula transcriptome, relative to control (c) (k=2, stress=0.08). **c,d,** MA plots of transcriptomes changes between 2h and 24 h (c) and 2h and 48 h (d). Blue dots denote genes with significant time-dependent expression changes regardless of baclofen treatment, red dots denote significant differences in time-dependent changes with baclofen treatment. Black dots denote genes with non-significant changes in expression. Negative log fold change values indicate inhibition.

### Autoregulation of GABA biosynthesis

Since baclofen is not metabolized in the cell in comparison to GABA, we postulated that the prolonged activation of the GABA_B_R pathway by baclofen would reduce the biosynthesis of GABA. Therefore, we tested for changes in expression of putative homologs that are relevant to this pathway. Indeed, we found that the expression of the glutamate transporter homologs EAAT1, EAAT3 and EAAT5, which in vertebrate carry the GABA substrate glutamate into the presynaptic cell, as well as the expression of four homologs of glutamic acid decarboxylases (GADs), which decarboxylate glutamate into GABA, were down regulated in planulae treated with baclofen (Fig. 6a,b). Furthermore, genes coding for enzymes that degraded GABA or reuptake GABA from the synaptic cleft, such as GABA-transaminase (GABA-T), GABA transporter 1 (GAT1) and inhibitory vesicle transporters (vGATs), were upregulated in the presence of baclofen. A second GABA transporter homolog, GAT2, was down regulated in baclofen-treated planulae. Interestingly, tumor susceptibility gene 101 (TSG101), which is a component of the ESCRT (endosomal sorting complex required for transport) complex and is involved in GABA_B_R degradation in vertebrates (44–46), was up regulated following baclofen addition and down regulated following baclofen wash. We also found that three GABA_A_ receptor subunits were down regulated after 48h baclofen treatment as well as one GABA_B_ (v1g239821) receptor. However, the expression level of all the other GABA_B_ subunits did not change. The finding of down-regulation of GABA synthesis pathways and up-regulation of GABA degradation pathways imply that GABA levels will be reduced in baclofen-treated planulae. Indeed, testing GABA spatial accumulation in control and baclofen-treated planulae demonstrated strong GABA reduction in treated planulae (Fig. 6c,d). These results suggest the presence of functional GABA signaling and of GABA regulatory pathways in *Nematostella*, where baclofen activates a GABA_B_R dependent signaling pathway and triggers a specific autoregulation response that includes inhibition of GABA synthesis and activation of GABA removal and degradation.

**Figure 6:**
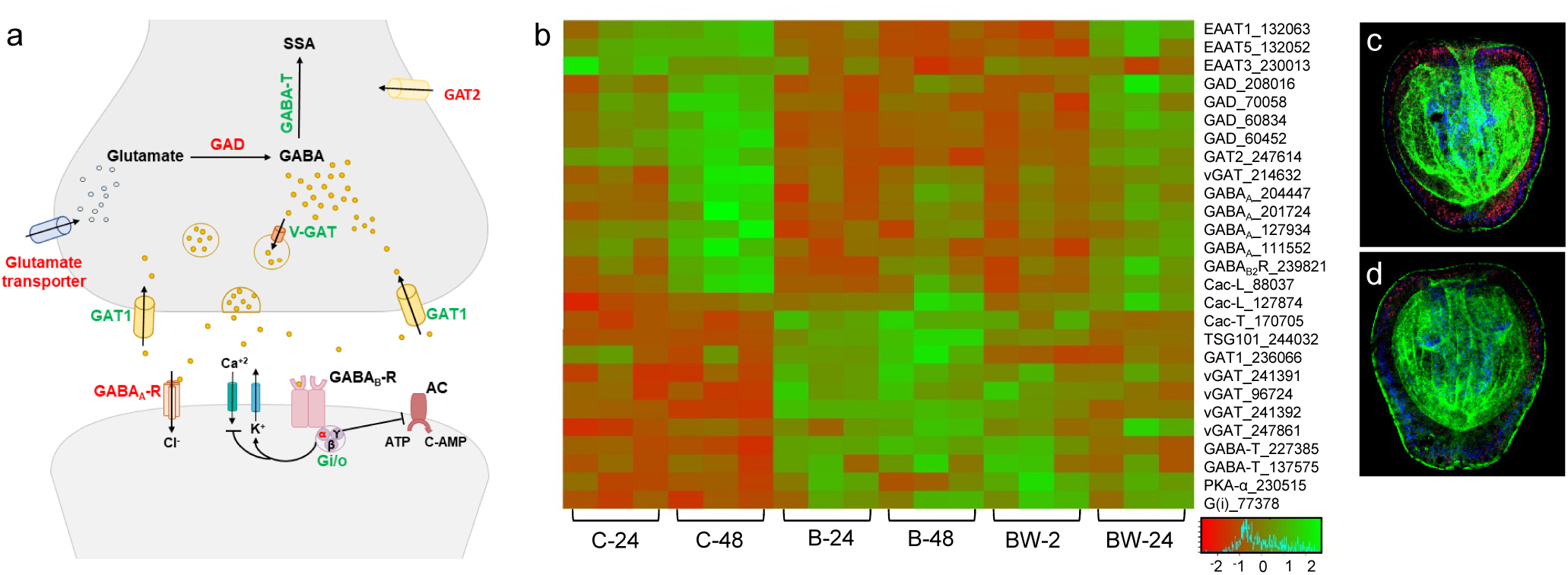
Autoregulation of GABA synthesis following baclofen treatment. **a,** Schematic illustration of the GABA pathway within the human synapse, based on (74). Transcripts that were up-regulated following baclofen treatment are colored green and transcripts that were down-regulated following baclofen treatment are colored red. **b,** Heat map of transcript expression associated with GABA bio-synthesis pathways as a function of time and treatment (C, control; B, baclofen treatment; BW, baclofen/wash). **c-d,** Confocal sections of control planulae (c) and 48-h baclofen-treated planulae (d) labeled with antibodies against GABA (red), phalloidin (green) and DAPI (blue).

### GABA signaling inhibits planula neurogenesis

Next, we examined the effect of prolonged GABA_B_R activation on early development of the nervous system of planulae. Transcriptomic analysis revealed significant down regulation in baclofen-treated planulae of pro-neural transcription factors such as *NvSoxB(2)*, *NvAshA*, *NvAth-like*, *NvNeuroD1*, *NvOtxB*, *NvSoxC*, and *NvBrn2*, of two neuron-specific RNA-binding proteins, *NvElav1* and *NvMsi*, and of other neuronal genes (Supplementary Table 1). Spatial expression analysis of key neurogenic transcription factors in control and baclofen-treated planulae demonstrated that while their expression levels decreased, their expression patterns did not change (Fig. 7a,b). An exception to this down regulation trend was *NvSoxB1* that initially, in the transcriptomic data, was down regulated in baclofen-treated planulae and then up regulated to a similar expression level as in the control (Fig. 7a,b).

**Figure 7:**
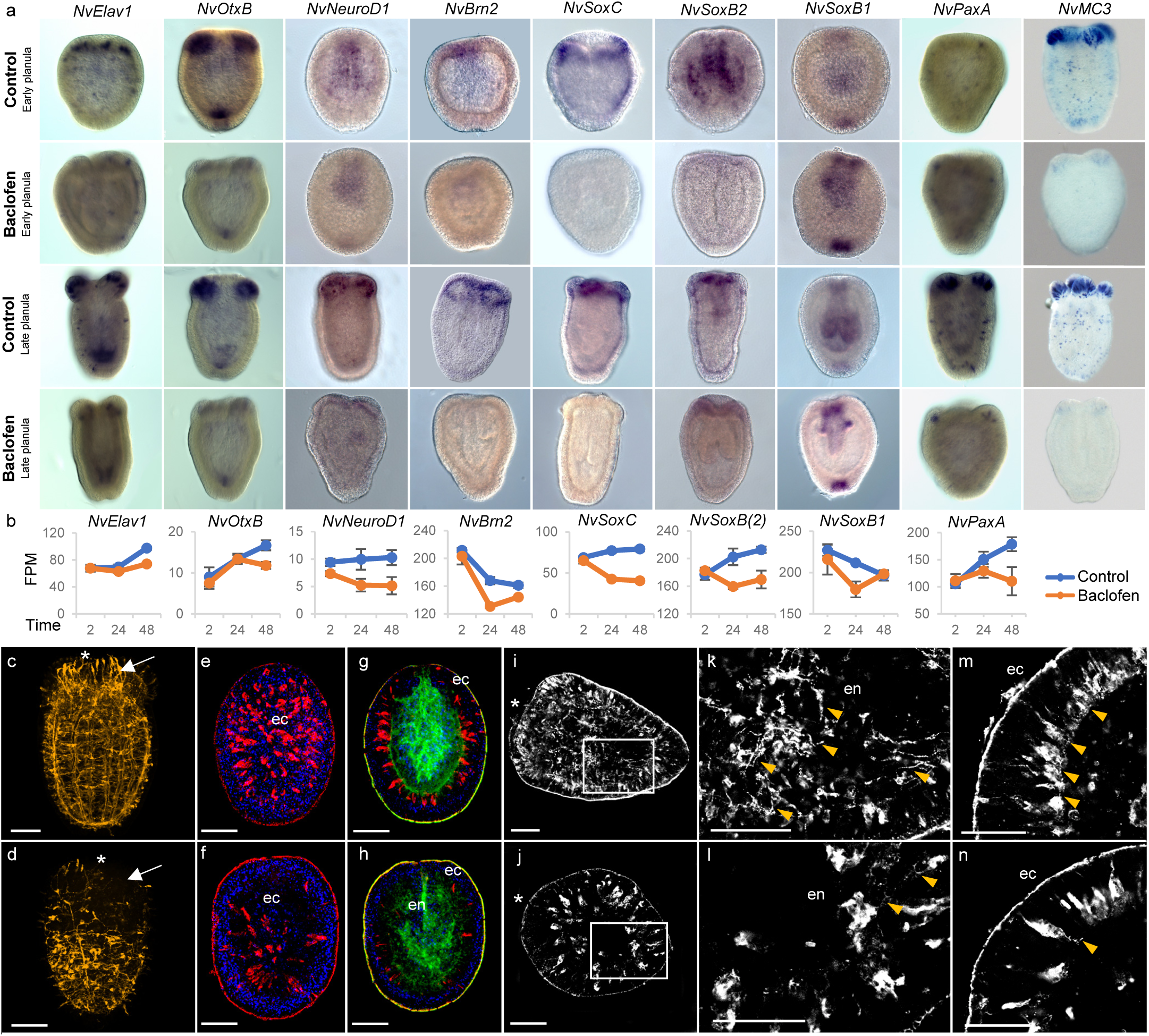
Baclofen inhibits *Nematostella* neurogenesis. **a,** Whole-mount in situ hybridization for neurogenesis-related genes in early and late control and baclofen-treated planulae. **b,** Transcriptome analysis of the significantly differentially expressed neural genes using FPM (fragments per million) 2, 24, 48 hours post baclofen addition and in control planulae. **c,d**, Confocal sections of 6dpf *NvElav1::mOrange* control (c) and baclofen-treated planulae (d) labelled with an anti-mCherry antibody. Note the less organized Elav-expressing neuron net in baclofen-treated planulae, specifically in the oral part (marked with arrows). **e-h** Confoca l sections of 4 dpf control planulae (e,g) and baclofen-treated planulae (f,h) labeled with antibodies against RF-amide (red), phalloidin (green) and DAPI (blue). Note that RF-amid neurons are organized in the ectoderm. **i-n**, Confocal sections of 6 dpf control planulae (i,k,m) and baclofen-treated planulae (j,l,n) labeled with anti-RF-amide antibody (white). The areas marked with white rectangles in i and j are magnified in k and l respectively. Note the long and developed neurites (marked with yellow arrows) in control planulae (k,m) versus baclofen treated-planulae (l,n), which have few and short neurites. Oral side is facing up in all images or labeled with asterisk. ec-ectoderm; en-endoderm; Scale bars, 50 µm.

We further tested a transgenic line expressing mOrange under the promotors of *NvElav1*. Addition of baclofen reduced *NvElav1*gene expression, resulting in a less organized neural net that lacked the longitudinal neurons along the mesenteries, forming a less dense mesh-like structure, specifically in the oral pole (Fig. 7c,d). In untreated planulae *Elav* positive neurons are elongated in the ectoderm of the oral pole creating a “crown like” structure that is absent in baclofen-treated planulae. The cnidarian nervous system also includes the stinging cells known as cnidocytes; therefore, we also analyzed genes specifically expressed during cnidogenesis (Fig. 7a). Expression of mini-collagen 3 (*NvMc3*), a structural protein in the stinging capsule, is extremely reduced in baclofen-treated planulae, as was expression of *NvPaxA*, a member of the homeodomain transcription factors family that is required for cnidocyte development (47).

The substantial down regulation of key positive regulators of neural differentiation following baclofen treatment indicated that prolonged activation of GABA_B_R signaling inhibits neurogenesis. Thus, we expected to find less differentiated neurons in baclofen-treated planulae. To test this hypothesis, we performed immunostaining with an antiRF-amide neuropeptide antibody. We compared untreated planulae at 4 and 5 dpf to planulae treated with baclofen for 24 h or 48 h. At early planula stages, RF-amide-positive neurons were found in the ectodermal layer (Fig. 7e-h). As the planula grows, these neurons form a net throughout the ectoderm and endoderm with high density in the oral region (Fig. 7i-n). We detected RF-amide neurons at the ectoderm layer in both control and baclofen-treated planulae. However, a significant decrease in these RF-amide neurons was observed 24 h after baclofen treatment (Fig. 7f,h). Following neural differentiation at 5 dpf, control planulae exhibited extended neurites connecting the neurons in both ectoderm and endoderm layers, as well as the presence of basiepithelial neurites that connect the ectodermal neurons (Fig. 7i,k,m). By contrast, in 5 dpf planulae following 48 h of baclofen treatment, only a few neurons developed neurites and these were relatively short. Additionally, only a few basiepithelial neurites were observed and no connections between RF-amide neurons in the ectoderm were seen (Fig. 7j,l,n). Hence, we conclude that the GABA_B_R signaling pathway inhibits neuron formation and nervous system development in *Nematostella*.

## Discussion

In this study, we demonstrated that activation of GABA_B_R by either a specific agonist or by a positive allosteric modulator inhibited planula-to-polyp transformation. This effect was reversible and showed no toxicity, while removal of GABA_B_R modulators led to a continuation of the *Nematostella* developmental plan. Interestingly, in other marine invertebrate such as molluscs and sea urchins, GABA was suggested to play a role in metamorphosis, either as an inducer or as an inhibitor, but receptors mediating these effects have not been characterized nor additional GABA signaling components have been implicated (48–50). In contrast, our characterization of putative *Nematostella* GABA_B_R homologs identified four genes with all the conserved features of a functional GABA_B1_R. The cysteines shown to form disulfide bridges that are essential for a functional GABA_B_R VFT extracellular domain (5) are conserved in all four putative GABA_B1_R *Nematostella* homologs, as were residues that are essential for a binding site for GABA and GABA agonists.

Our analysis also suggested that *Nematostella* GABA_B1_R proteins have smaller GABA binding pockets that will not accommodate the larger GABA_B_R antagonists. Indeed, we found that the latter had no effect on *Nematostella* planula-to-polyp transformation. Interestingly, in the cnidarian freshwater hydra, GABA_B_R agonists and antagonists were shown to affect the feeding response, but receptors responsible for this phenotype have not been characterized (51, 52). However, such differences in the ability of GABA_B1_ receptors to bind agonists or antagonists were also found in other species, despite the high conservation of previously characterized GABA_B1_R homologs among vertebrates and invertebrates. For example, baclofen had no effect in *D. melanogaster* and in *C. elegance*, and highly effective antagonists for mammalian GABA_B1_R had no effect in *Drosophila* (19, 53).

In addition to the identification of GABA_B1_R homologs, we also showed a functional connection to broader GABA regulation pathways. Baclofen affected the expression of diverse enzymes in GABA metabolism, reducing GABA biosynthesis and transport and increasing GABA removal and degradation – suggesting a functional autoregulated GABA signaling system in *Nematostella.* Since we found that several GABA_A_ receptors were also down regulated following baclofen application, this supports cross-talk between GABA_B_ and GABA_A_ signaling, as was suggested before in mammalian systems (54).

As the main inhibitory neurotransmitter in metazoans, GABA is an essential part of the mammalian nervous system. It was demonstrated that in the mammalian brain, activation of GABA_B_R promote quiescence of neural stem cells and inhibition of neurogenesis (13, 16, 17). Our results suggest an evolutionarily conserved function in *Nematostella*, as baclofen activation of GABA_B_R inhibited expression of pro-neuronal genes and neurogenesis. Additionally, there was a significant decrease in the Elav- and RF-amide neuron population and in the development of neurites in baclofen-treated planulae. We also found that genes related to cnidocyst synthesis and regulation were down-regulated following baclofen treatment, which is in line with previous finding in hydra, suggesting that GABA_B_R play a role in cnidocyst discharge (51). However, unlike the sustained transcriptomic down regulation of neuron related genes after baclofen addition, *NvSoxB1* expression quickly recovered to a level similar to control planulae. In Bilateria, the SoxB family is divided into two subgroups: SoxB1 and SoxB2, which have antagonizing functions – controlling neuron stem cell maintenance and neuron differentiation (55). In vertebrates, *SoxB1* genes are expressed in most progenitor stem cells of the developing nervous system and in adult neurogenic regions, and their activity maintains cells as proliferative precursors and prevents neural differentiation (56–58). Conversely, *SoxB2* expression counteracts and represses *SoxB1* activity and promotes neural differentiation (59). In *Nematostella*, the exact classification of SoxB subgroups is not clear (60–62). While not much is known on *NvSoxB1* (63, 64), *NvSoxB(2)* was shown to be required for the development of neural progenitor cells, including cnidocytes (65, 66). Our results support the latter findings, as baclofen caused down regulation of *NvSoxB(2)*, downstream genes involved in neurogenesis, as well as in cnidocytes-synthesis related genes.

Adding to previous studies, which positioned *Nematostella* as a model system for evolutionary comparisons and developmental biology, our study highlights *Nematostella* as a new model system for GABA signaling. Thus, our results open up new avenues for comparative studies of GABA signaling function and evolution. Furthermore, the rapid and simple developmental cycle of *Nematostella* and the ease of high-throughput screening of modulators establish *Nematostella* as an exciting and simple model organism for further exploration of the extended GABA signaling pathway.

## Methods

### Sea anemone culture

Anemones were cultured in *Nematostella* medium (NM) composed of 12.5 ppt artificial sea water (Red Sea, Israel) maintained at 18°C in the dark. Anemones were fed 5 times a week with freshly hatched *Artemia* brine shrimps. Mature Sea anemones were induced to spawn as described (67). Embryos were raised at 21°C in the dark and planulae or polyps were collected for the different experiments.

### Pharmaceutical treatments

Treatments were done in triplicates, with each plate containing approximately 100 planulae. GABA_B_R agonists: GABA (10^−3^ M, 10^−4^ M, Sigma-Aldrich), baclofen (10^−4^ M, Sigma-Aldrich), or CGP-7930 (10^−5^ M, Sigma-Aldrich) (35) were added at 1, 3 or 4 dpf. Baclofen and GABA were dissolved in water, using NM as a control, whereas CGP-7930 was dissolved in DMSO, using NM with 10^−5^ M DMSO as a control. Planulae were raised in the dark at 21°C. To remove the agonists the medium was replaced with NM and washed 5 times. To test metamorphosis rate, primary polyps were counted each day. The effect of GABA_B_R antagonists on metamorphosis was tested using Phaclofen (10^−3^ M, 10^−4^ M, Sigma Aldrich) and the high affinity antagonist CGP-54626 (10^−7^ M, 10^−6^ M, 10^−5^ M, Cayman). Muscimol (10^−4^ M, Sigma Aldrich) was used to test whether GABA_A_ receptor play a role in metamorphosis.

### Identification and cloning of *Nematostella* GABA_B_ receptors

To uncover *Nematostella* homologs of human GABA_B_R we searched with the sequences of human GABA_B1b_R (NP068703) and GABA_B2_R (CAA09942) using blastp all *Nematostella* proteins in the NCBI RefSeq database. Putative identified homologs were used as queries against the UniProt database to confirm homology. In addition, we also searched for *Nematostella* homologs with the N-terminal extracellular region and the transmembrane regions of the human queries separately, using both the blastp and tblastn options. The following four *Nematostella* full sequences were identified: v1g244104, v1g239821, v1g206093, v1g243252. Full length sequences of the three previously identified partial *Nematostella* GABA_B1_R homologs (v1g86565, v1g158857, v1g87697) and a newly identified homolog (v1g210496) were PCR-amplified using the following primers (with NCBI identifiers): 5’, ATGTCAAGTGTCGGAGCTATTG and 3’, TCACTCTTTTGATTGCATCGGAC for NvGABA_B1a_ (MH194577); 5’, AGACCAAAGGCCGACTCACAA and 3’, TGACAAACCGATATACCGCGA for NvGABA_B1b_ (MH194578); 5’, ATGTTCATTAATTTCTTGTGGCCTGT and 3’, CCTTAGTTAATAAATTTATTTGCGAGA for NvGABA_B1c_ (MH194579); 5’, CAGAATGAACTGGCACAAGC and 3’, ACGCATGCAAAAATACAATATCTTTT for NvGABA_B1d_ (MH355581), followed by sequencing.

### Domain predictions and three-dimensional structural analysis

*Nematostella* putative GABA_B_R genes were translated to proteins using the ExPASy online translate tool (http://web.expasy.org/translate). Sequence conservation was visualized using the ExPASy online boxshade tool (http://www.ch.embnet.org/software/BOX_form.html). Conserved domains in *Nematostella* GABA_B_R proteins were identified using the consensus of three different databases: the NCBI Conserved Domain Database (https://www.ncbi.nlm.nih.gov/cdd), InterPro (http://www.ebi.ac.uk/interpro) and SMART (http://smart.embl-heidelberg.de). To identify predicted transmembrane regions, we used a consensus among the servers TMHMM (http://www.cbs.dtu.dk/services/TMHMM), TOPCONS (http://topcons.net) and SignalP4.1 (http://www.cbs.dtu.dk/services/SignalP). Coiled-coil domains were predicted using a consensus of SMART (http://smart.embl-heidelberg.de), pcoils (https://toolkit.tuebingen.mpg.de/#/tools/pcoils) and Paircoil2-MIT (http://cb.csail.mit.edu/cb/paircoil2).

We generated 3D models of the extracellular regions of NvGABA_B_ proteins using Swiss-model (https://swissmodel.expasy.org) (68) using the crystal structure of the extracellular domain of human GABA_B_R – PDB ID 4MS4 (4). The following protein data bank (PDB) crystal structures were used: 4MS3, structure of the extracellular domain of human GABA_B_R bound to the endogenues agonist GABA; 4MS4, structure of the extracellular domain of human GABA_B_R bound to the agonist baclofen; 4MR7, structure of the extracellular domain of human GABA_B_R bound to the antagonist CGP54626. Visualization of the 3D models was done using the PyMOL Molecular Graphics System (https://pymol.org).

### RNA extraction, sequencing and bioinformatics analysis

Transcriptome experiments were done on 4 dpf baclofen-treated (10^−4^ M) and control planulae in triplicates of 200 planulae per sample. Samples were taken for RNA extraction from controls and after baclofen addition at 2 h, 24 h and 48 h. In addition, samples of planulae treated with baclofen for 24 h were washed, and collected 2 h and 24 h after wash. Samples were frozen in TriReagent (Sigma-Aldrich) at ^-^80°C. Total RNA was extracted using TriReagent (Sigma-Aldrich) according to the manufacturer’s instructions. The RNA was further purified using the RNA Clean & Concentrator TM-5 kit (Zymo Research), and genomic DNA residues were removed by DNase treatment (Ambion). RNA quality and concentration were determined using an Agilent 2200 TapeStation.

The 24 samples were prepared for multiplex sequencing using the NEB Ultra Directional RNA kit according to the manufacturer’s instructions. Samples were sequenced using 50-bp single-end reads in two lanes on an Illumina HiSeq2000 and a TruSeq v3 flow chamber at the Life Sciences and Engineering Infrastructure Center at the Technion, Haifa. Illumina reads were quality-filtered and adapter-trimmed using Trimmomatic, and inspected with Fastqc (www.bioinformatics.babraham.ac.uk). Filtered reads (about 12-18 million reads, for the different samples) were mapped and quantitated using the *Nematostella vectensis* genome (NCBI genome assembly GCA_000209225.1 ASM20922v1), using STAR (version 2.4.2a) (69). 72%-75% of all reads were mapped uniquely, and ∼15% were mapped to multiple loci. Differential expression analysis was conducted using DESeq2 (70). Transcript abundances were represented as FPKMs (fragments per kilobase million), and significance was defined as adjusted p-value < 0.05. Expected factor-dependent trends were verified using non-metric multidimensional scaling (NMDS) graphs in Vegan (https://github.com/vegandevs/vegan). Heatmaps were generated using shinyheatmap (http://shinyheatmap.com) (71)based on Ward’s hierarchical clustering, considering FPKM-based Euclidean distances between genes.

### *In situ* hybridization and immunohistochemistry

Whole-mount in situ hybridization (ISH) was done according to (72) and immunohistochemistry was performed according to (73). The following primary antibodies were used: anti-GABA (1:500, abcam ab86186), anti-mCherry (to detect mOrange) (1:200, Abcam ab167453) and anti-FMRFamide (Merck Millipore AB15348). The following secondary antibodies were used: Donkey anti-mouse Alexa Fluor 488 (Jackson 715-545-150), Goat anti-rabbit rhodamine (Jackson 711-295-152) and Goat anti-rabbit Alexa Fluor 647 (Jackson 111-605-144). F-actin labeling was done using Alexa Fluor 488-conjugated phalloidin (2 u/ml 66 nM, Sigma-Aldrich) and nucleus staining was done with DAPI (1:1000, Sigma-Aldrich).

### Microscopy and imaging

*In situ* hybridization was visualized using a Zeiss Axio Imager 2 epifluorescence microscope equipped with an AxioCam MRm camera (Carl Zeiss, Jena, Germany). Immunofluorescent was visualized using confocal microscopy (Nikon, C1-SHS UMT). Planulae were visualized using a Nikon binocular (Multizoom Az-100) or Nikon eclipse Ti.

## Acknowledgements

We thank F. Rentzsch for providing the Elav reporter line. We thank the Bioinformatics Service Unit at University of Haifa and specifically M. Lalzar and A. Malik for their assistance. This research was supported by the Israel Ministry of Science and Technology (grant 3-8774) and the Israel Science Foundation (grants 1454/13, 2155/15).

## Author Contributions

S.L. designed and performed the experiments, V.B. performed gene cloning and assisted in the experiments, S.L., A.B., and M.K. performed sequence and structure analysis, A.S-P analyzed GABA_B_R homologs expression in the single cell dataset, S.L, V.B., M.K., and T.L. analyzed the data, M.K. supervised sequence and structure analysis, T.L. conceived and supervised the project, T.L wrote the manuscript with input from all authors. All authors discussed the results and commented on the manuscript. All authors read and approved the final version of the manuscript.

## Competing interest statement

The authors declare no competing financial interests.

---

**Supplementary Figure 1:**
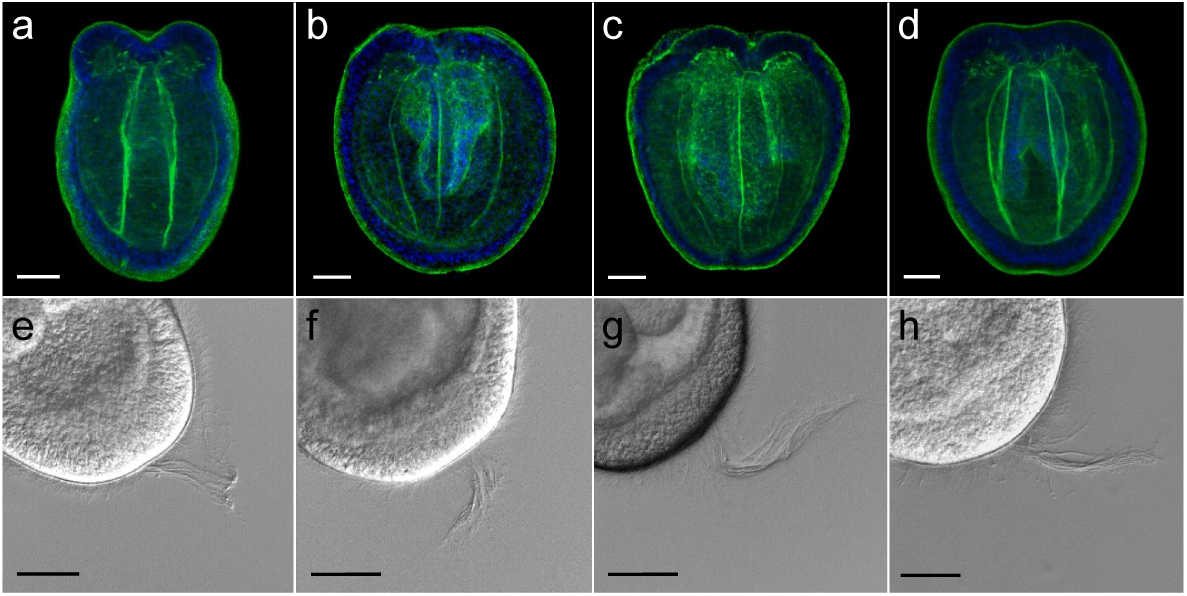
Morphological comparison of control and GABA_B_R modulators. **a-d,** Confocal sections of control and treated planulae labeled with antibodies against phalloidin (green) and DAPI (blue); **e-h** DIC pictures of the apical tuft; Control planulae (a,e), treated planulae with GABA (10^−3^ M) (b,f), baclofen (10^−4^ M) (c,g) and CGP-7930 (10^−4^ M), a GABA_B2_R positive allosteric modulator (d,h); Scale bars, 50 µm.

**Supplementary Figure 2:**
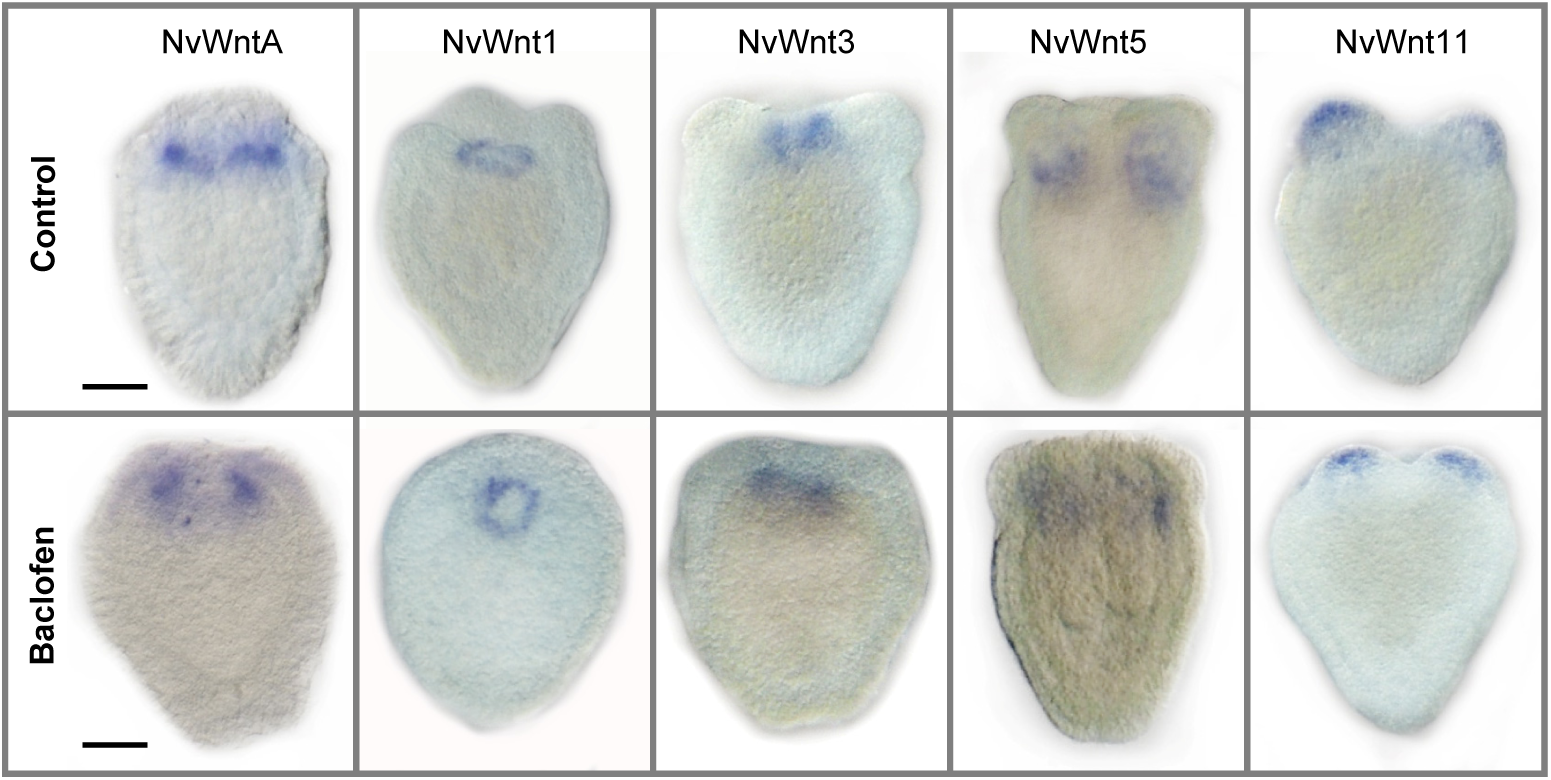
GABA_B_R activation does not affect the Wnt ligands pattern expression. In situ hybridization for Wnt ligands of control and 48-h baclofen-treated planulae. Expression levels and patterns of WntA (v1g91822), Wnt1 (v1g158342), Wnt3 (v1g241352), Wnt5 (v1g235248), and Wnt11 (v1g230011) did not change following baclofen treatment; Scale bar, 50 µm.

**Supplementary Figure 3:**
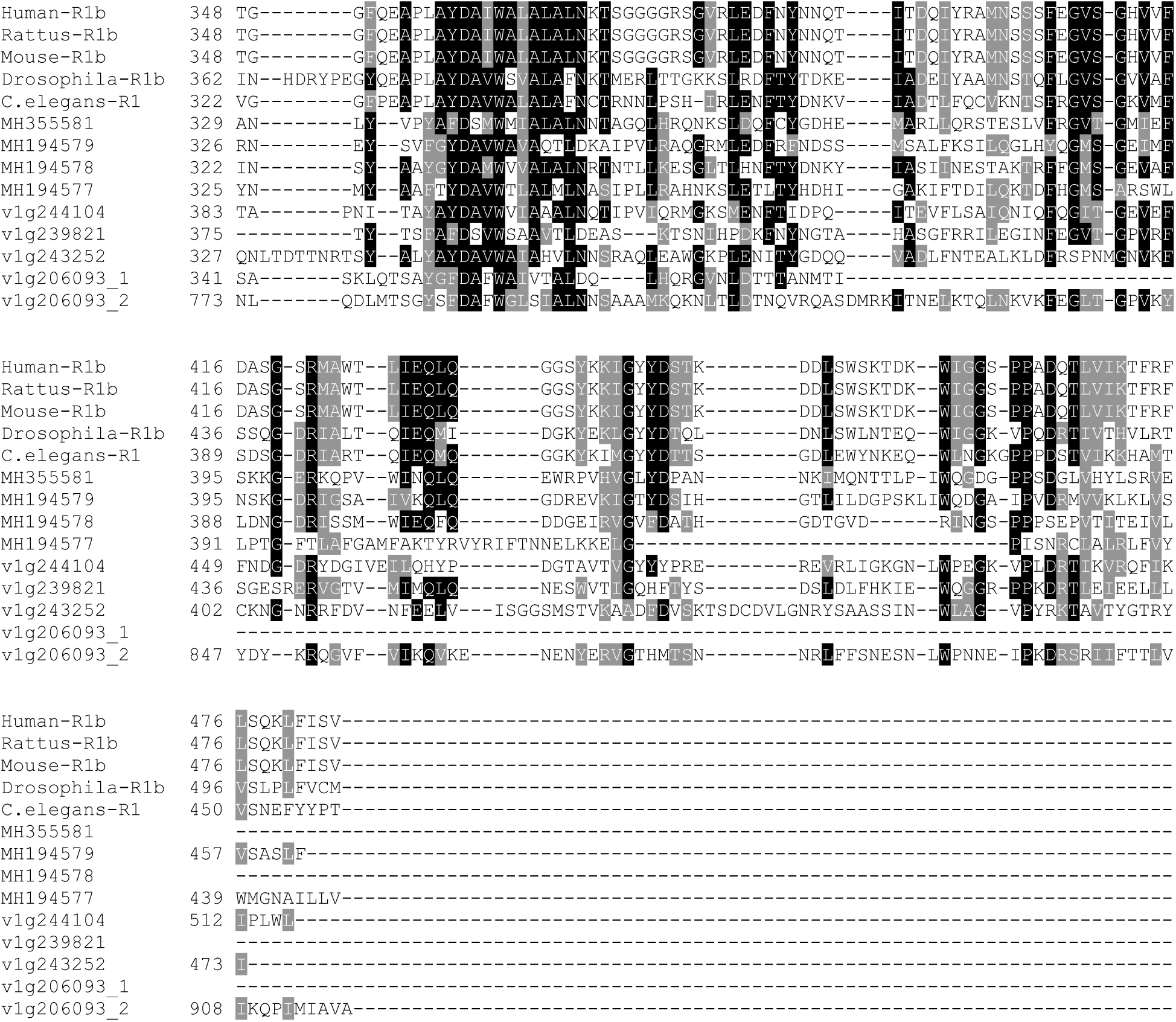
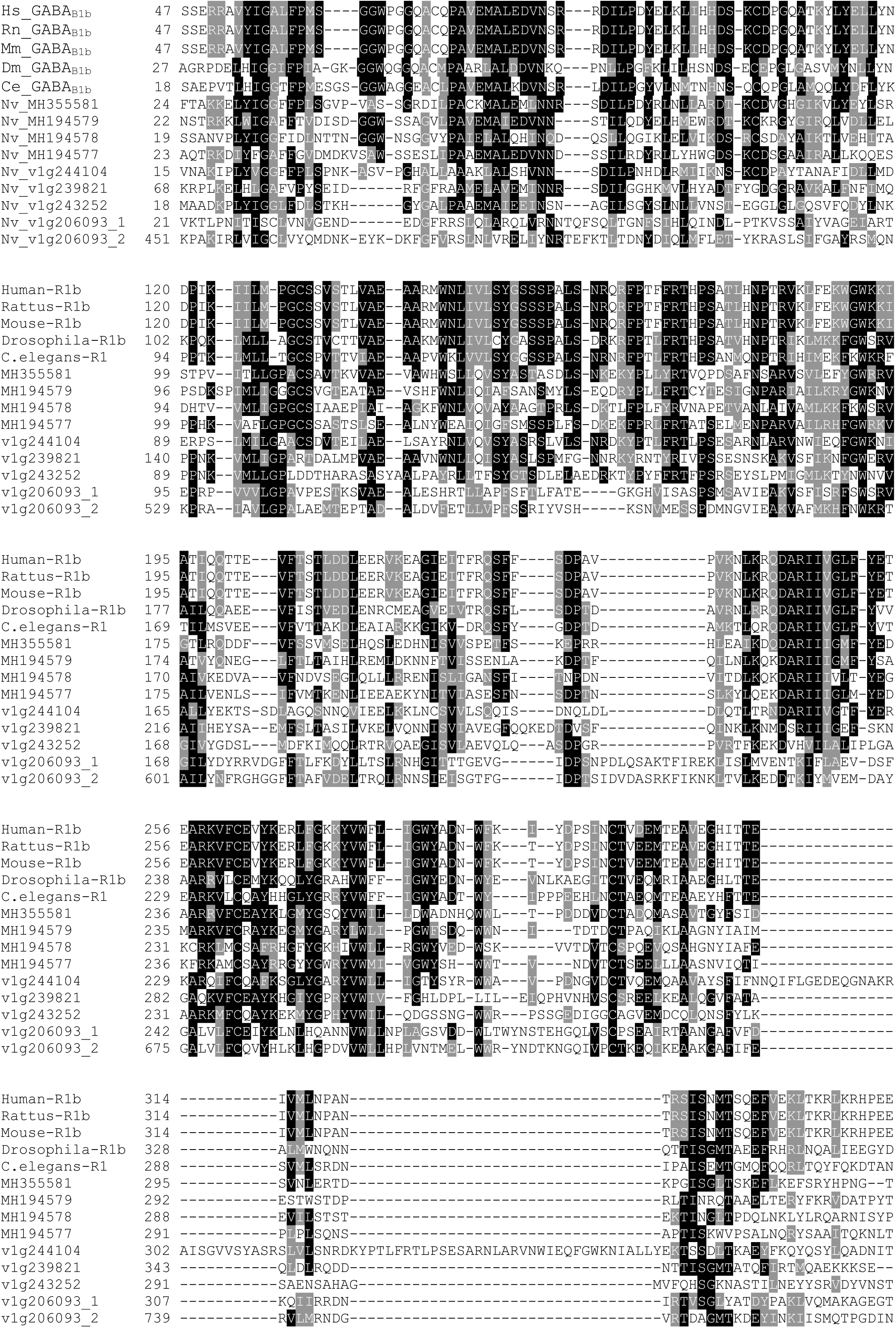
Alignment of the extracellular region of bilaterian GABA_B1_R homologs and the eight putative *Nematostella* GABA_B_R homologs. Identical residues are marked in black and conservatively substituted residues are marked in gray. Bilaterian sequences are as follows, with NCBI Genebank accession numbers: human (Hs), NP_068703; rat (Rn), AAD19657; mouse (Mn), AAH56990; *Drosophila* (Dm), AAF53431; *C. elegans* (Ce), ACE63490, followed by the eight identified *Nematostella* (Nv) GABA_B_-R homologs. The two extracellular domains of *Nematostella* v1g206093 are shown separately.

**Supplementary Figure 4:**
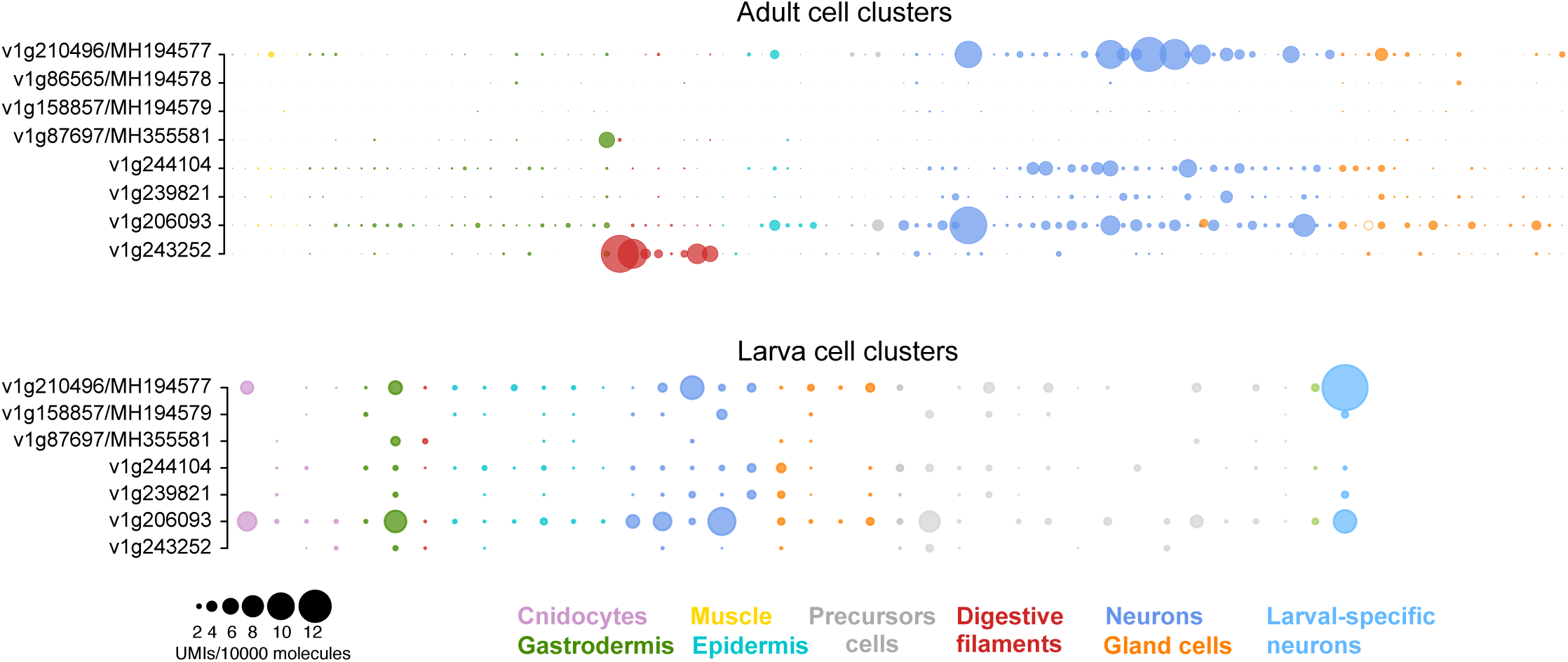
Expression of GABA_B_R homologs across cell clusters from the *Nematostella* adult and larva single-cell atlas (43). Gene expression levels are computed as Unique Molecule Identifiers (UMIs) per every 10,000 samples UMIs in each cell cluster.

